# Dendritic spines on GABAergic neurons respond to cholinergic signaling in the *Caenorhabditis elegans* motor circuit

**DOI:** 10.1101/598714

**Authors:** Andrea Cuentas-Condori, Ben Mulcahy, Siwei He, Sierra Palumbos, Mei Zhen, David M. Miller

## Abstract

Dendritic spines are specialized postsynaptic structures that detect and integrate presynaptic signals. The shape and number of dendritic spines are regulated by neural activity and correlated with learning and memory. Most studies of spine function have focused on the mammalian nervous system. However, spine-like protrusions have been previously reported in invertebrates, suggesting that the experimental advantages of smaller model organisms could be exploited to study the biology of dendritic spines. Here, we document the presence of dendritic spines in *Caenorhabditis elegans* motor neurons. We used super-resolution microscopy, electron microscopy, live-cell imaging and genetic manipulation to show that GABAergic motor neurons display functional dendritic spines. Our analysis revealed salient features of dendritic spines: (1) A key role for the actin cytoskeleton in spine morphogenesis; (2) Postsynaptic receptor complexes at the tips of spines in close proximity to presynaptic active zones; (3) Localized postsynaptic calcium transients evoked by presynaptic activity; (4) The presence of endoplasmic reticulum and ribosomes; (5) The regulation of spine density by presynaptic activity. These studies provide a solid foundation for a new experimental paradigm that exploits the power of *C. elegans* genetics and live-cell imaging for fundamental studies of dendritic spine morphogenesis and function.

**HIGHLIGHTS:**

- Spines in *C. elegans* GABAergic motor neurons are enriched in actin cytoskeleton.
- Spines are dynamic structures.
- Spines display Ca^++^ transients coupled with presynaptic activation.
- Spine density is regulated during development and is modulated by actin dynamics and cholinergic signaling.

## INTRODUCTION

The majority of excitatory synapses in the mammalian brain feature short, local protrusions from postsynaptic dendrites that respond to presynaptic neurotransmitter release^1^. These dendritic “spines” were originally described by Ramon y Cajal^2^ and are now recognized as key functional components of neural circuits. For example, spine morphology and density are regulated by neural activity in plastic responses that are strongly correlated with learning and memory^3, 4^. Although spine-like protrusions have been reported for invertebrate neurons^5, 6^, few studies^7, 8^ have rigorously determined if these structures share functional features with vertebrate spines.

The anatomy of the *C. elegans* nervous system was originally defined by reconstruction of electron micrographs (EM) of serial sections. Although this approach revealed that most *C. elegans* neurons display smooth dendritic surfaces, a subset of neurons was also reported to display short, spine-like protrusions. These include five classes of motor neurons (RMD, RME, SMD, DD, VD) and an interneuron (RIP)^9, 10^. Subsequent work using light and electron microscopy detected similar dendritic protrusions extending from D-class motor neurons in the related nematode, *Ascaris*^11, 12^. Finally, a recent report used light microscopy to show that a postsynaptic acetylcholine receptor is localized near the tips of spine-like protrusions on DD class motor neurons that directly appose presynaptic termini^13, 14^. Here, we have adopted a systematic approach to test the hypothesis that these spine-like structures in *C. elegans* GABAergic motor neurons (DD and VD) exhibit the salient hallmarks of dendritic spines in mammalian neurons.

## RESULTS

### Dendritic spines in *C. elegans* GABAergic neurons

In the adult, Dorsal D (DD) class GABAergic motor neurons extend axons to innervate dorsal muscles and receive cholinergic input at ventral dendrites (Figure 1A). We used AiryScan imaging, a type of super-resolution microscopy^15^, to detect spine-like projections on the ventral processes of adult DD neurons labeled with a cytosolic mCherry marker (Figure 1B). Because the actin cytoskeleton is a structural hallmark of vertebrate dendritic spines^16^, we also labeled DD neurons with the actin marker, LifeAct::GFP^17^. Super-resolution images detected apparent enrichment of LifeAct::GFP in DD spines versus the dendritic shaft. For a quantitative assessment, we calculated the ratio of the spine to shaft fluorescence and plotted the cumulative distribution for each marker (Figure 1E). This representation shows a clear separation between measurements of cytosolic mCherry that is evenly distributed throughout dendrites (median spine/shaft ratio < 1) versus that of the LifeAct::GFP signal (median spine/shaft ratio > 1) (KS test, p<0.0001, Figure 1E). Thus, actin is enriched in DD spines. This interpretation is strengthened by our finding that independent measurements of the spine density (mean spine numbers/10 µm) with either cytoplasmic mCherry or LifeAct::GFP are not significantly different. Similar results were obtained by EM reconstruction (see below) (Figure S1A).

**Figure 1.**
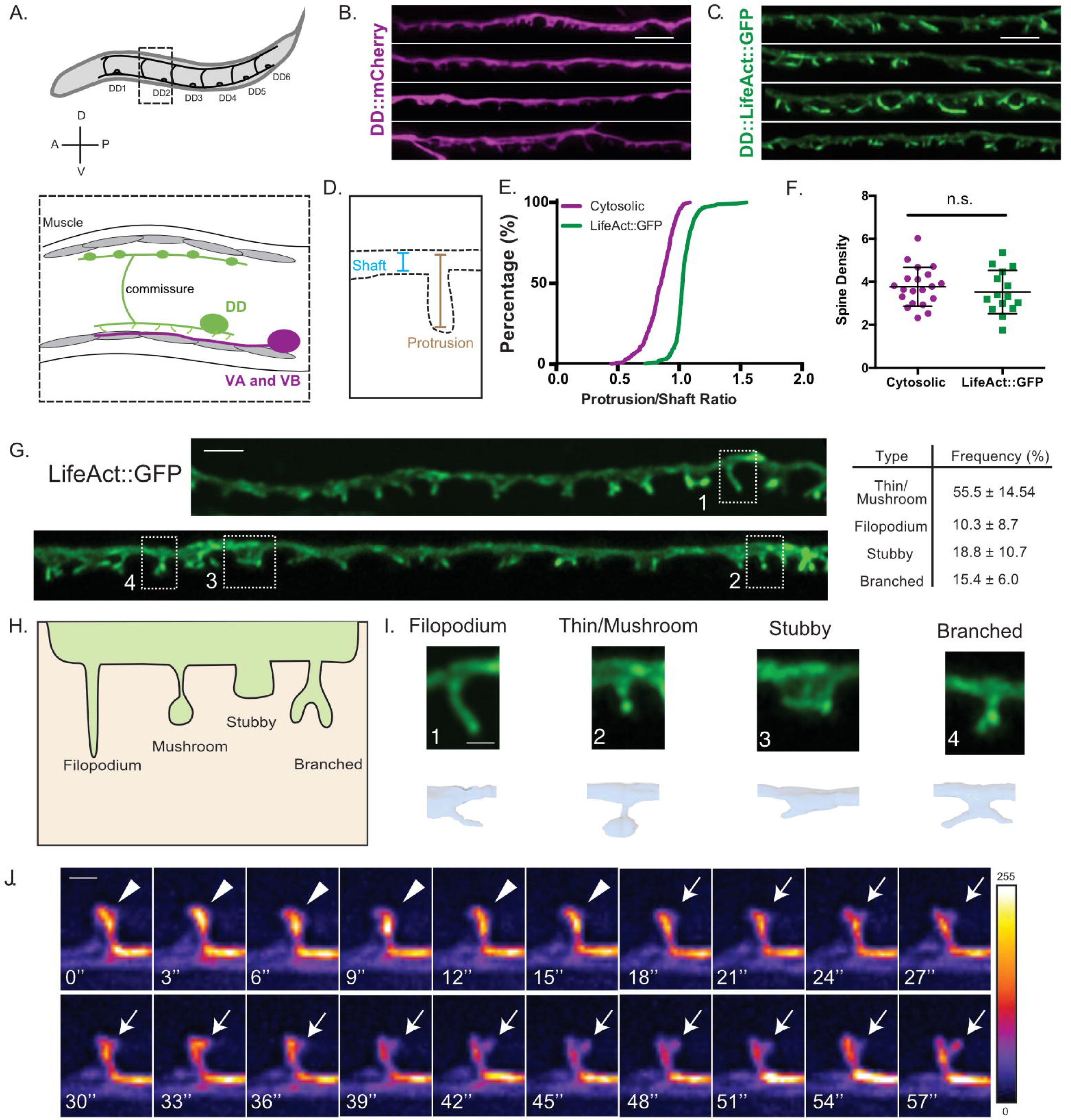
DD GABAergic neurons display dendritic spines. **A.** Six DD motor neurons are located in the *C. elegans* ventral nerve cord. In the adult, DD presynaptic boutons (oblong ovals) innervate dorsal muscles (grey cells) and DD postsynaptic termini (spines) receive cholinergic input from VA and VB motor neurons on the ventral side (purple). **B-F.** AiryScan imaging resolves ventrally projecting spine-like protrusions from DD neurons labeled with **(B)** cytosolic mCherry or **(C)** LifeAct::GFP. **(D)** Intensity of protrusion over shaft ratio reveals that **(E)** LifeAct::GFP preferentially accumulates at the protrusion whereas cytosolic mCherry is evenly distributed between the protrusion and shaft (KS test, p<0.0001, n > 286 spines). **(F)** Spine density of young adults revealed by mCherry (3.8 ± 0.9 spines/10µm) or LifeAct::GFP (4.3 ± 1.1 spines/10µm) is not significantly different (t test, p=0.0855, n > 16 worms). Measurements are mean ± SD. Scale bars = 2 µm. **G-I.** LifeAct::GFP reveals 1. Thin/Mushroom (55.54 ± 14.54%), 2. Filopodial (10.27 ± 8.70%), 3. Stubby (18.78 ± 10.74%), 4. Branched spines (15.42 ± 6.01%) in adult DD motor neurons. Measurements are mean ± SD, n = 16 worms. For scatterplot, see Figure S1. (Scale bar = 1 µm). **(H)** Schematic of spine shapes. **(I)** Images of each type of spine identified by (top) AiryScan imaging (Scale bar = 500nm) of LifeAct::GFP or (bottom) 3D-reconstruction of DD1 from serial electron micrographs of a high-pressure frozen adult. See also Figure S1B. **J.** Dendritic spines are dynamic. Snapshots of *in vivo* spine remodeling from a thin (arrowhead) to branched morphology (arrow). Images (LifeAct::GFP) are shown with rainbow LUT. Higher intensity is represented by warm colors and dimmer intensity by cold colors. L4 stage worm. See also Figure S1C-H. Scale bar = 500nm

A close inspection of LifeAct::GFP-labeled protrusions revealed a variety of spine shapes which we grouped into several morphological classes resembling those previously reported for mammalian dendritic spines^1, 18^: thin/mushroom, filopodial, stubby and branched (Figure 1G-I) (see Methods). We merged “thin” and “mushroom” shapes into a single category because both are defined by an enlarged head region vs a narrower neck. By these criteria, adult DD neurons have predominantly thin/mushroom spines, with lesser fractions of filopodial, stubby and branched shapes (Figure 1G).

As an independent method for assessing the presence of dendritic spines, we used EM to reconstruct the anterior-most 25 µm dendrite of DD1, the most anterior member of the DD class of motor neurons^10^. For this experiment, young adult animals were prepared using High Pressure Freezing (HPF) to avoid potential artifacts arising from chemical fixation^10, 19^. Reconstruction of 50 nm serial ultrathin sections from the anterior DD1 dendrite detected twelve DD1 spines, with multiple morphological shapes that resemble classes revealed by LifeAct::GFP labeling (Figure 1i; Figure S1B). Our findings are consistent with observations from EM reconstructions of mammalian dendritic spines in the hippocampus where thin and mushroom shapes predominate ^20, 21^ and filopodial and stubby spines are less abundant^22, 23^.

### DD spines are shaped by a dynamic actin cytoskeleton

Although we have assigned DD motor neuron spines to four discrete classifications, both LifeAct::GFP imaging and EM reconstruction point to a broader array of spine types that includes potential intermediate forms (Figure 1G-H). A similarly heterogeneous array of spine shapes among mammalian neurons has been attributed to active remodeling of spine architecture^23, 24^.

To test for this possibility in *C. elegans*, we used live imaging to produce time-lapse recordings of DD spines. Our live-imaging revealed that some DD spines can remodel *in vivo* (Figure 1J). During imaging sessions of ≥ 30 minutes, we observed several cases (11 out of 25 movies) of transient filopodial-like extensions from the dendritic shaft that destabilize in the course of minutes (See Supplementary Video S1-S4). Most DD spines, regardless of the type, were stable throughout a given imaging session. In the mature mammalian cortex, extended imaging has revealed transient filopodial extensions with a lifetime shorter than a day^23, 25^, and potential plasticity over longer intervals, where approximately half of spines are stable for months^23, 25^. To our knowledge, our time lapse images are the first to directly visualize dynamic dendritic spines in a motor neuron of a living organism^26^.

Live-imaging of DD motor neurons also detected a dynamic actin cytoskeleton (Figure S1C-G), consistent with previous reports for mammalian dendritic spines^27, 28^. To ask if actin assembly is required for DD spine morphogenesis^16^, we applied genetic methods to knockdown key regulators of actin polymerization and assessed their effect on DD spines (Figure S1H). We found that the Arp2/3 complex, and two of its activators, the F-BAR protein TOCA-1 and the Wave Regulatory Complex, are required to maintain DD spine density (Figure S1H). Disruption of the Arp2/3 complex or its activators has been previously shown to reduce dendritic spine density in the mammalian brain and to impair function^29–33^.

### Dendritic spines of DD neurons directly appose presynaptic terminals

Because functional dendritic spines are sites of presynaptic input^5, 34, 35^, we investigated the disposition of DD spines vis-a-vis their main presynaptic partners, the cholinergic VA and VB class motor neurons (Figure 2A). For super-resolution imaging, we used the synaptic vesicle-associated marker, mCherry::RAB-3, to label the presynaptic termini of VA and VB class cholinergic motor neurons in the ventral cord and LifeAct::GFP to label DD neurites and spines (Figure 2B). Clusters of mCherry::RAB-3-labeled puncta are located adjacent to DD spines (Figure 2B)^13^. Among the 128 spines identified by LifeAct::GFP, most (∼84 %) reside near at least one presynaptic cluster (denoted “contacted” in Figure 2D)^13^. Approximately ∼40% (51/128) of DD spines are associated with multiple presynaptic clusters of RAB-3 puncta which suggests that individual spines receive input from > 1 presynaptic terminal (arrowheads, Figure 2B).

**Figure 2.**
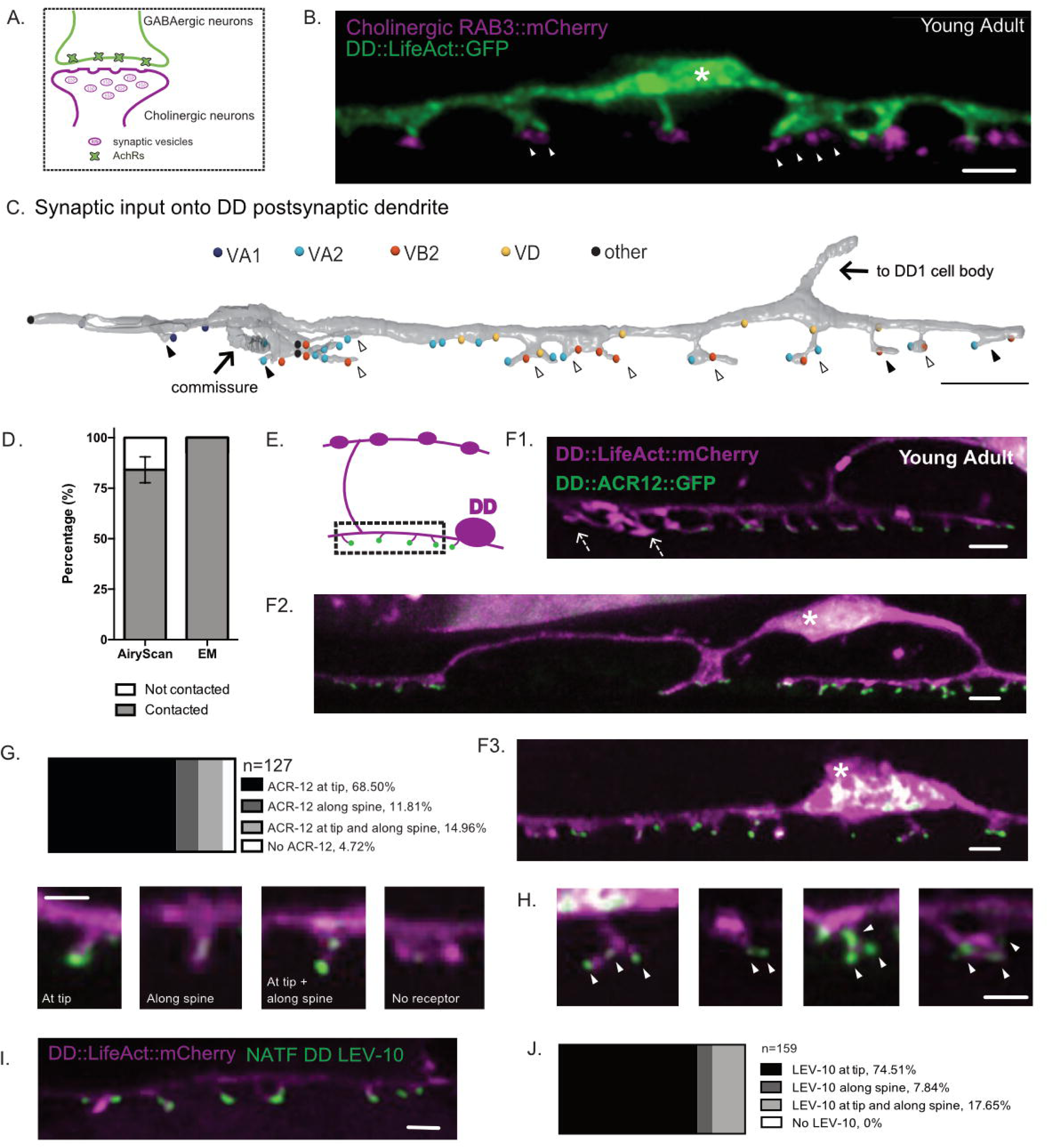
DD spines appose presynaptic cholinergic vesicles. **A.** Ionotrophic acetylcholine receptors (iAChRs) are localized in GABAergic motor neurons in apposition to input from cholinergic motor neurons. **B.** Cholinergic RAB-3 presynaptic vesicles labeled with mCherry localize in close proximity to DD postsynaptic spines labeled with *flp-13::*LifeAct::GFP. Scale bar = 1µm. Asterisk denotes the DD cell body. Arrowheads mark multiple RAB-3::mCherry clusters apposing dendritic spines. **C.** Volumetric EM reconstruction of a portion of the DD1 dendrite (25 µm) in the ventral nerve cord (gray) detects contacts with 43 different presynaptic termini, from axons of the cholinergic (VA1, VA2, VB2) and GABAergic (VD) motor neurons, and other neurons (other). 84.8% (n=28/33) of VA and VB inputs are adjacent to DD spines. 33.3% (4/12) of spines directly oppose a single presynaptic partner (black arrowhead); the majority (66.67%) appose more than one terminal (clear arrowhead). Scale bar = 2 µm. **D.** Schematic of DD presynaptic boutons (top) and postsynaptic spines (dashed box) with distal AChR puncta (green dot) on the ventral side. **E1-2.** ACh receptor subunit ACR-12::GFP (green) localizes to LifeAct::mCherry-labeled DD spines (magenta). Asterisk marks DD cell body. Arrows in E2 denote spines without ACR-12::GFP clusters. Scale bars = 1 µm. **F-G.** Locations of ACR-12::GFP puncta on DD spines. > 95% of spines have at least one ACR-12::GFP cluster (n = 127 spines from 8 worms). **(G)** Examples of spines from each category. Scale bars = 1 µm. **H.** Examples of spines with more than one ACR-12::GFP cluster. White arrowheads point to ACR-12::GFP cluster at DD spine. **I-J.** NATF labeling of endogenous AChR auxiliary protein LEV-10 in DD neurons (I) detects LEV-10 localization to spines with **(J)** all spines showing NATF LEV-10::GFP puncta (n = 159 spines from 7 worms). Scale bars = 500nm.

Our EM reconstruction revealed direct apposition of all DD spines (n = 12/12) with the presynaptic termini of cholinergic motor neurons (Figure 2C). Further analysis of the DD1 dendrite and cholinergic motor neuron (VA1, VA2 and VB2) axons in this region showed that most (84.8%, n = 28/33) cholinergic presynaptic inputs appose DD1 spines, whereas a smaller number (15.2%, n = 5/33) are positioned along the dendritic shaft. This finding parallels the observation that only 10% of excitatory synapses in the mature mammalian cortex are positioned on dendritic processes^16^. Two thirds of DD1 spines (n = 8/12) receive input from more than one neuron. That is, a single DD1 spine head is contacted by presynaptic termini of both VA and VB class cholinergic motor neurons (Figure 2C and Video S5). This finding could explain the observation above from Airyscan imaging that multiple clusters of RAB-3 puncta are adjacent to ∼40% of DD spines (Figure 2B). We note that individual spines on GABAergic neurons in the mammalian hippocampus can also have input from multiple presynaptic termini^5, 36, 37^. DD1 spines receive some inhibitory synapses from the other class of GABAergic motor neurons (VDs). Most of these VD inputs are restricted to the DD1 dendritic shaft (n = 5/6) (Figure 2C).

The acetylcholine receptor (AChR) subunit ACR-12 is postsynaptic to cholinergic inputs at GABAergic motor neurons, and has been previously shown to localize to DD1 dendritic protrusions (Figure 2E)^13, 38, 39^. We used AiryScan imaging to quantify the subcellular distribution of the ACR-12::GFP signal on DD spines. We detected ACR12::GFP clusters on ∼95% of DD spines (n = 121/127) (Figure 2E), with 68.5% (n = 87/127) localized at spine tips and the remainder either positioned along the lateral side of the spine (∼12%, n = 15/127) or at both the side and tip (∼15 %, n = 19/127) (Figure 2G). 43.3% of spines (n = 55/127) had more than one ACR-12::GFP cluster (Figure 2H), a finding that mirrors the recent observation that the spines of mammalian cortical neurons can display multiple postsynaptic assemblages of the postsynaptic protein PSD-95^40^.

To obviate the possibility that the localization of ACR-12::GFP clusters to DD spines results from marker over-expression, we used a new live-cell labeling scheme to detect an endogenous component of the acetylcholine receptor complex, the co-factor protein LEV-10^41^. NATF (Native and Tissue-Specific Fluorescence) relies on the reconstitution of superfolder GFP (sfGFP) from the split-sfGFP fragments, GFP1-10 and GFP11^42^. We used genome editing to fuse seven tandem copies of GFP11 to the C-terminus of the native LEV-10 coding sequence. GFP1-10 was then selectively expressed in DD neurons from a transgenic array (i.e., *flp-13::GFP1-10*). In this case, we detected LEV-10 NATF-GFP signal at 100% (n = 159/159) of DD spines (Figure 2I-J), further substantiating the hypothesis that DD spines are sites of presynaptic input.

### ER and ribosomes localize to DD neuron dendritic spines

Key cytoplasmic organelles such as Smooth Endoplasmic Reticulum (SER), have been reported in both the dendritic shaft and spines of mammalian neurons^21^. In addition to its role of processing membrane proteins, the SER in spines is known to regulate activity-dependent Ca^++^ release through the ryanodine receptor^43^. Other structures such as polysomes and rough ER have also been reported in spines, raising the possibility of local translation^18, 44, 45^.

Our EM reconstruction of DD1 revealed cisternae SER-like structures in both the dendritic shaft and spines (Figure 3A-B). Ribosome-like structures were detected in some DD1 spines (Figure 3B”). This observation is consistent with our independent finding that the ribosomal protein, RPS-18::GFP^46^ is also detected in DD spines (Figure 3C). Mitochondria and microtubules are reported to be rare in the dendritic spines of mature mammalian neurons^18, 44^. Our EM reconstruction did not reveal mitochondria or microtubules in all twelve DD spines (Figure 3D). Both organelles were detected in DD1 dendritic shaft (Video S6). These observations however should be interpreted cautiously given the small number of spines reconstructed.

**Figure 3.**
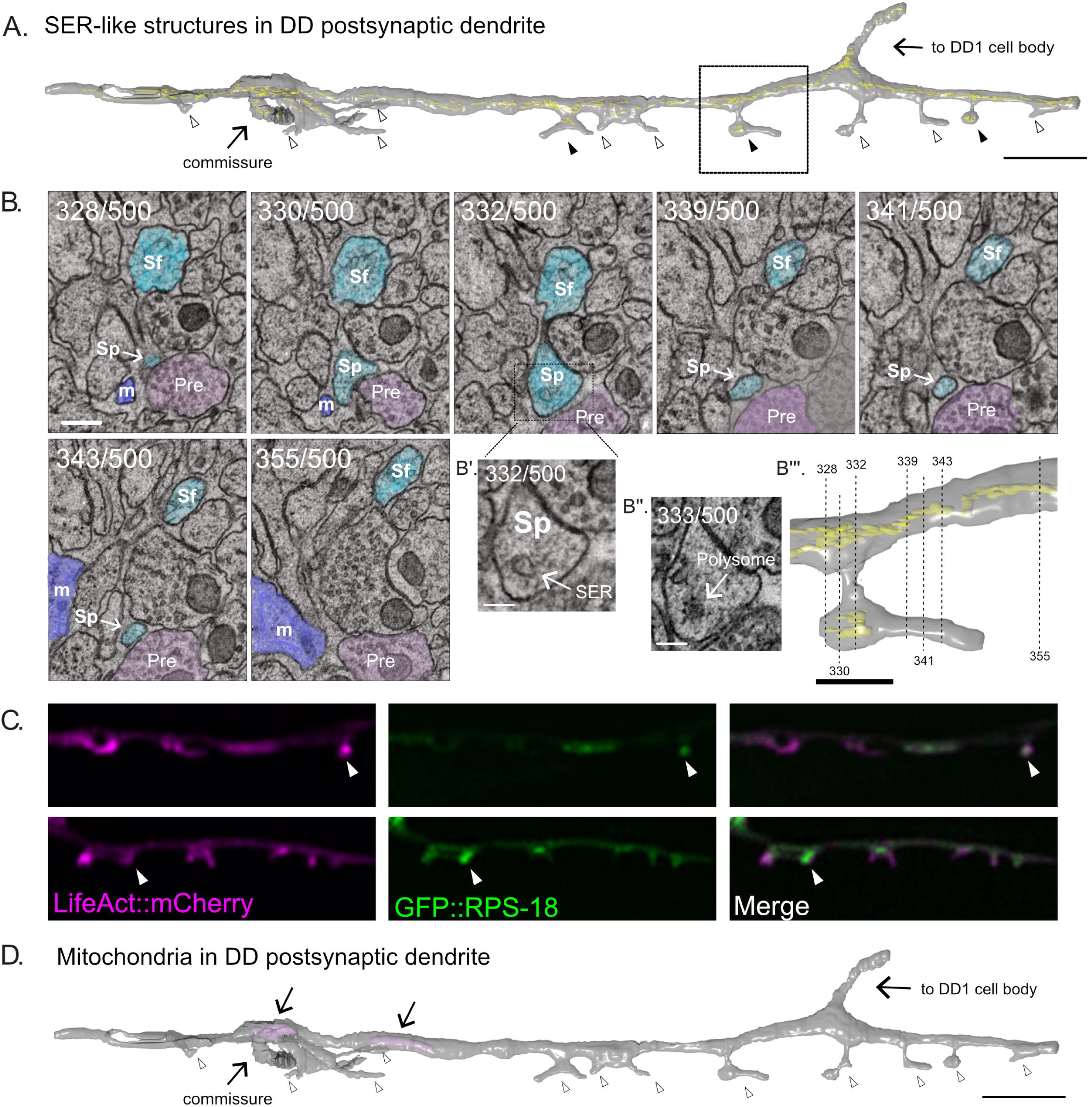
SER-like structures and ribosomes localize in spines and dendritic shaft. **A.** 3D EM reconstruction of DD1 dendrite reveals SER-like cisternae (yellow) in the dendritic shaft and some spines (black arrowheads). Most spines lack SER-like structures (clear arrowheads). Scale bar = 2 µm. **B.** Serial cross-sections (328 – 355) of the ventral nerve cord show spines (Sp) budding from DD1 (blue) dendritic shaft (Sf). ‘Pre’ labels presynaptic terminals from a cholinergic VA neuron (pink); m, muscle arm (purple). (Scale bar = 200 nm) **B’.** Magnified region of section 332. Arrow points to SER-like structure in DD dendritic spine (Scale bar = 100 nm). **B”.** Section 333. Arrow points to polysome-like structure in DD dendritic spine (Scale bar = 100 nm). **B”’.** Volumetric reconstruction of DD1 dendrite (gray) and SER-like structures (yellow). Dashed lines denote location of each section shown in B. Scale bar = 500 nm. **C.** AiryScan imaging shows GFP-labeled ribosomal protein, RPS-18^46^, localized to DD spines (arrowheads) labeled with LifeAct::mCherry. **D.** Volumetric EM reconstruction of the portion of the DD1 (25 µm) dendrite shows mitochondria (purple) in the shaft (arrows) but not in spines (clear arrowheads).

### Activation of presynaptic cholinergic motor neurons drives Ca^++^ transients in DD spines

Ca^++^ is one of the main signaling molecules to mediate activity-dependent synaptic plasticity^1, 47^. We reasoned that if DD spines are functional, we should detect dynamic Ca^++^ transients. To test this hypothesis, we expressed the Ca^++^ sensor GCaMP6s in DD neurons. Live-imaging (at 1 second intervals) revealed spontaneous Ca^++^ transients in both DD spines and shafts (Figure 4A-C and Video S7-S8). Interestingly, Ca^++^ transients were observed simultaneously in adjacent spines about 50% of the time (Figure 4D). To estimate the likelihood of simultaneous Ca^++^ peaks in adjacent spines occurring by chance, we compared the distribution of the observed time differences between neighboring spine Ca^++^ peaks (△T) to a uniform distribution of time differences (at 1 second intervals) over the period of observation (10 seconds). These distributions are statistically different (KS test, p<0.0001), which suggests that the observation of correlated Ca^++^ dynamics may reflect mechanisms for linking postsynaptic activity in adjacent DD spines (Figure 4D). Simultaneous Ca^++^ dynamics among neighboring spines has been reported for developing hippocampal neurons in culture. It will be interesting to determine if adjacent spines on mature mammalian motor neurons are also coordinately activated^26^. In these cases, correlated Ca^++^ waves between neighboring spines showed long decay times and were mainly driven by intracellular calcium release from the SER after initial glutamate receptor activation^47^. Consistent with this idea, SER-like structures were observed in both DD1 dendritic shaft and spines (Figure 3A-B).

**Figure 4.**
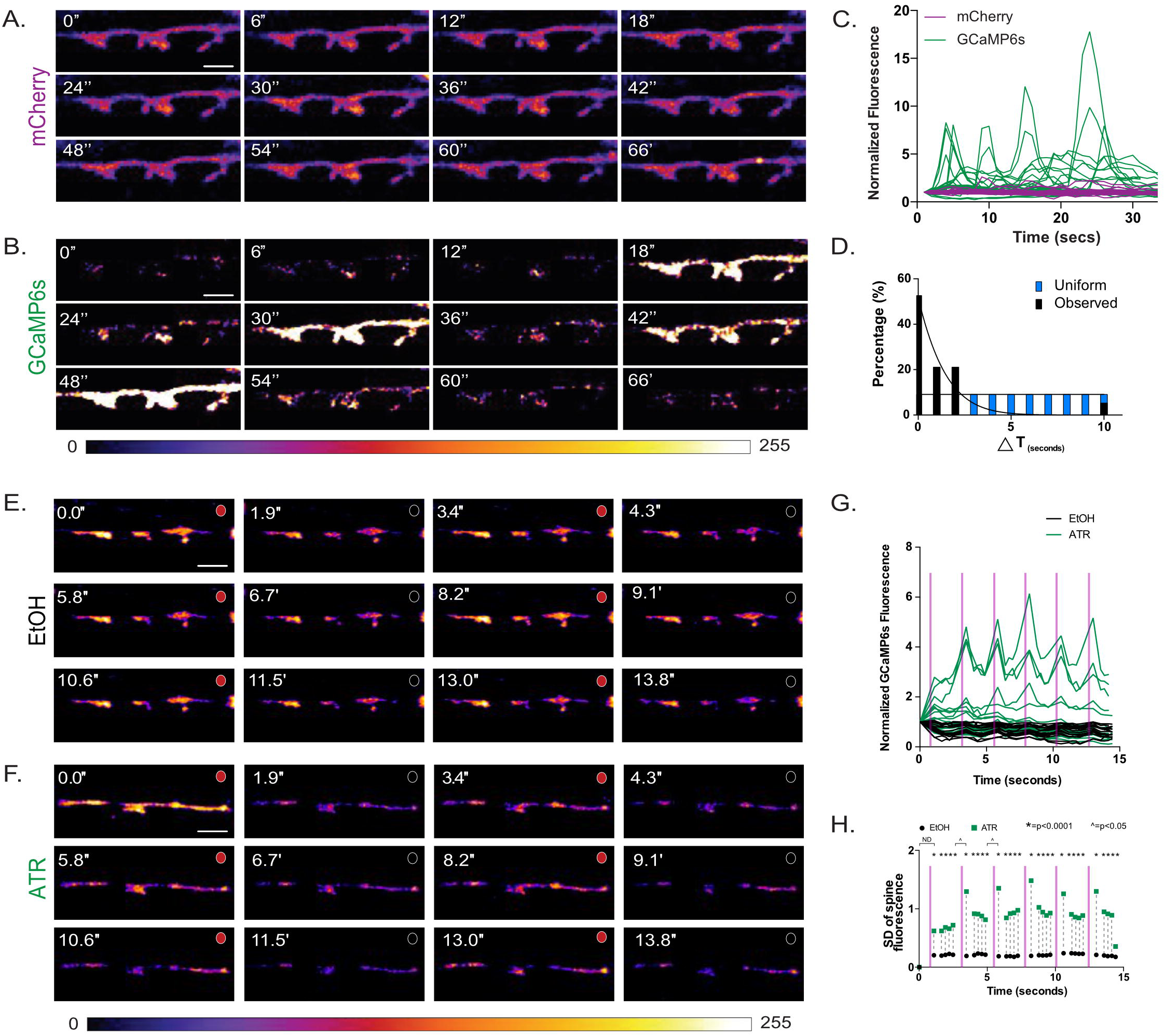
Ca^++^ transients in dendritic spines. Series (time in minutes) of live-cell images of cytosolic **(A)** mCherry and **(B)** GCaMP6s in DD postsynaptic spines reveals (**C**) dynamic GCaMP6s vs stable mCherry signals, n = 11 movies, 31 spines. **D.** GCaMP6s transients occur in neighboring spines more frequently (> 50%) than predicted by a random distribution (KS test, p<0.0001). Scale bars = 500 nm. **E-H.** VA motor neuron activation is correlated with Ca^++^ transients in DD1 spines. GCaMP6s fluorescence imaged (at 0.5 sec intervals) with periodic (at 2.5 sec intervals) optogenetic activation of ceChrimson, detects Ca^++^ transients with (**E**) ATR (n = 14) but not with carrier (EtOH) (**F**) (n = 12). Circles at the top of each panel correspond to red light on (red) for ceChrimson activation vs off (black). Scale bars = 500 nm. (**G**) GCaMP6s fluorescence throughout the 15 sec recording period plotted for ATR (green) (n = 14) vs carrier (EtOH) (black) (n = 12). (**H**) Plot of the standard deviation (SD) of GCaMP6s signal at each time-point shows that fluctuations in the ATR-treated samples (green boxes) are significantly greater than in EtOH controls (black circles), F-test, *= p < 0.0001. Additionally, SDs are significantly different between timepoints before and after light activation (T_6_ vs T_7_ and T_11_ vs. T_12_). F-test, ^ = p < 0.05. ND = not determined. Purple bars denote interval with red-light illumination (e.g., ceChrimson activation).

We did not observe Ca^++^ signals in DD spines when cholinergic receptors were desensitized by administration of a cholinergic agonist levamisole (data not shown). These findings are consistent with the idea that Ca*^++^* transients in DD spines depend on presynaptic cholinergic signaling^13^. To test this idea, we engineered a transgenic animal for optogenetic activation of VA neurons with red-light illumination (*Punc-4:*:ceChrimson::SL2::3xNLS::GFP)^48^ and detection of Ca^++^ changes in DD spines with blue-light excitation (*Pflp-13*::GCaMP6s::SL2::mCherry). VA motor neurons are presynaptic to DDs and therefore are predicted to evoke DD neuronal activity^10, 13^. ceChrimson was activated by a brief flash of red light (80ms) at 2.5 second intervals and the GCaMP6s signal in DD1 spines was recorded at 2 Hz. This experiment detected a striking correlation of a GCaMP6s fluorescence after ceChrimson activation (Figure 4E-G and Video S9-S10). Although GCaMP6s fluorescence also varied in DD spines in the absence of red-light illumination, fluctuations were strongly correlated with ceChrimson activation as shown by a plot of the standard deviation of GCaMP6s fluorescence for all traces across the 15 second sampling period (Figure 4H). These results are consistent with the interpretation that DD spines are responding to cholinergic input from presynaptic VA motor neurons.

### Cholinergic signaling enhances DD spine density during development

In mammalian neurons, dendritic spine shape and density are modulated throughout development^20, 22, 49^. To determine if DD spine morphogenesis is also developmentally-regulated, we used the LifeAct::GFP marker to quantify spine density in four successive larval stages: L3, early L4, mid-L4 and young adult. This experiment revealed that both spine number and density increase as DD neurons elongate during development (Figure 5B and 5E).

**Figure 5.**
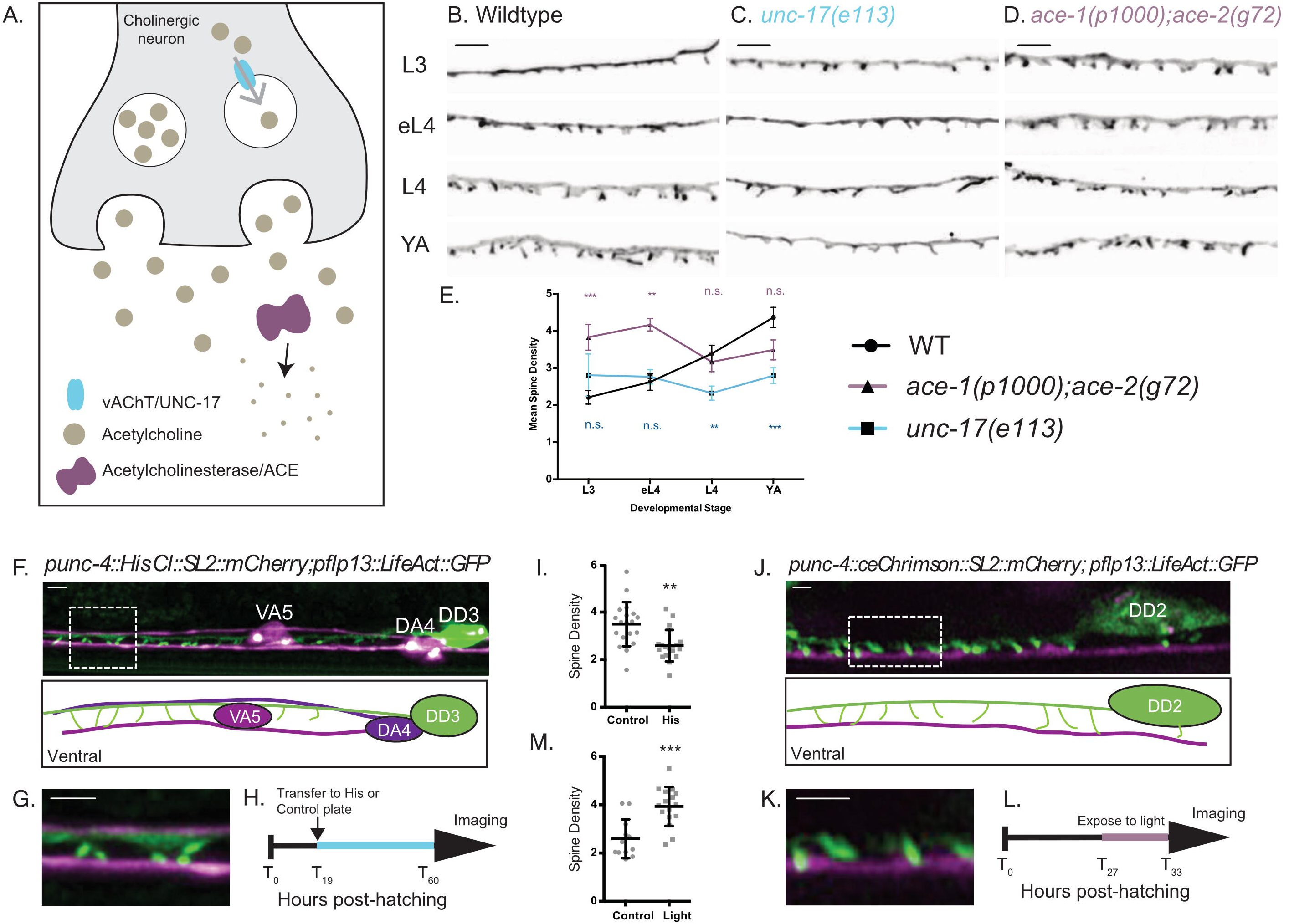
Cholinergic activity regulates spine density during development. **A.** Synaptic vesicles are loaded with acetylcholine (ACh) by the vesicular acetylcholine transporter (*unc-17*). Acetylcholinesterase enzymes (*ace-1 and ace-2*) degrade synaptic ACh. **B-E.** Spine density increases throughout development in the wild type (WT), but not in *unc-17(e113)* mutants whereas spine density is precociously elevated in *ace-1(p1000);ace-2(g72)* mutants. Representative images of **(B)** WT, **(C)** *unc-17 (e113)* and **(D)** *ace-1(p100); ace-2(g72)*. Scale bars = 2 µm. See Figure S2 for scatter plots for E. **F-I.** Reduced ACh signaling in cholinergic motor neurons decreases postsynaptic spine density. **(F)** Expression of Histamine-gated Chloride channels and mCherry in cholinergic (VA and DA) motor neurons (*punc4::HisCl::SL2::mCherry*^54^) vs DD motor neurons labeled with LifeAct::GFP shows **(G)** DD spines (green) extending to the ventral process of the VA5 motor neuron (purple). Note dorsal placement of DA4 axon. **(H)** Synchronized L1 larvae were transferred to either histamine or control plates at T_10_ (hours after hatching) for growth until adulthood (∼T_50_). See Methods. **(I)**. DD spine density is reduced by growth on histamine (2.58 ± 0.6) vs control (3.49 ± 0.9). T-test, ** = p<0.01, n > 17. Scale bars = 1 µm. **J-M**. Temporal activation of A-class cholinergic motor neurons increases spine density. **(J)** Cholinergic motor neurons (e.g., VA4) express ceChrimson^48^ and mCherry (*punc4::ceChrimson*^48^*::SL2::mCherry)*. LifeAct::GFP marks DD2. (**K**) Note DD spines (green) extending ventrally toward VA process (purple). **(L)** Synchronized L2 stage larvae (T_17_, hours post-hatching) were transferred to ATR plates (see Methods) for 5 hours (until T_23_) and exposed to red-light pulses vs control group grown in the dark. **(M)**. Exposure to red-light for 5 hours elevates spine density (3.9 ± 0.8) vs control (2.59 ± 0.8). T-test, *** is p<0.001, n>11. Scale bars = 1µm. See also Figure S2.

Dendritic spines, as sites of synaptic input, can be modulated by changes in synaptic strength^1, 18, 44^. Long-term potentiation, for example, is correlated with increased numbers of spines in the mammalian brain^25, 50, 51^. Similarly, hyperactivity is associated with increased dendritic spine density in hypoglossal motor neurons^26^. Conversely, long-term depression has been shown to induce spine shrinkage^52^ that may lead to their elimination. To test the idea that DD spines may also respond to changes in the strength of cholinergic signaling, we altered acetylcholine levels and assessed possible effects on spine density.

To reduce acetylcholine signaling, we used *unc-17/vAChT (e113)* mutants, in which expression of the vesicular acetylcholine transporter (UNC-17/vAChT) is selectively eliminated in cholinergic motor neurons (J. Rand, personal communication). For the converse condition of elevated acetylcholine signaling, we used mutants with reduced acetylcholinesterase activity, *ace-1(p100)* and *ace-2(g720)* (Figure 5A). Reduced synaptic acetylcholine release (i.e., in *unc-17/vAChT* mutant) results in lower DD spine density in the adult (Figure 5C and 5E). In contrast, increased synaptic acetylcholine (i.e., in *ace-1; ace-2* double mutants) results in precocious elevation of spine density during development (Figure 5D and 5E). These findings are consistent with the idea that cholinergic signaling positively regulates the formation of DD spines. The developmental elevation of spine density is also blocked in *unc-31/CAPS* mutants^53^, in which release of neuropeptide and catecholamine neurotransmitters is selectively prevented (Figure S2).

For an additional test of activity-dependent regulation of DD spine density, we modulated presynaptic cholinergic function for specific periods during larval development. To reduce cholinergic activity, we expressed the histamine-gated chloride channel^54^ in A class cholinergic motor neurons, which are direct presynaptic partners of DD neurons. Animals grown in the presence of histamine showed reduced spine density at the L4 stage compared to animals grown on plates without histamine (Figure 5F-I). To elevate cholinergic activity, we expressed the red-light activated opsin, Chrimson, in A-class motor neurons^48^. Animals were exposed to red light to activate Chrimson for a brief period (1 second every 4 seconds) during the L2-L3 stage larval development (See Methods). This treatment led to increased spine density (scored in L3) when compared to animals grown in the absence of red light (Figure 5J-M). These results demonstrate that DD spine density depends on presynaptic cholinergic signaling, thus confirming that DD spines share the fundamental property of mammalian dendritic spines of positive regulation by neuronal activity.

### VD-class GABAergic neurons also display dendritic spines

In the *C. elegans* motor circuit, dendrites of the DD-class GABAergic motor neurons receive cholinergic inputs in the ventral nerve cord, whereas the VD class dendrites are located in the dorsal nerve cord (Figure S3A). Because the original EM reconstruction of the *C. elegans* nervous system detected spine-like structures on VD neurons^9^, we sought to verify this finding by using the LifeAct::GFP marker for AiryScan imaging. We used miniSOG^55^ for selective ablation of DDs since LifeAct::GFP was expressed in both DD and VD neurons in this case and would otherwise obscure VD morphology. This experiment confirmed the presence of dendritic spines in VD neurons throughout the dorsal nerve cord (Figure S3A-D).

Our EM reconstruction of 27 µm of the anterior VD2 dendrite detected 9 dendritic spines (Figure S3E). Similar to DD1, most VD spines are closely apposed to presynaptic termini of cholinergic motor neurons (DA2, DB1, AS2). Additional presynaptic inputs from other cholinergic and GABAergic motor neurons (DD) are distributed along the dendritic shaft (Figure S3F) and several mitochondria are also observed in VD2 dendritic shaft (Figure S3E). Thus, our results confirm that both the DD and VD classes of ventral cord GABAergic motor neurons display dendritic spines^9, 10^.

## DISCUSSION

Although most studies of spine morphogenesis and function have been conducted in mammalian neurons, the prevalence of spine-like postsynaptic protrusions in invertebrate nervous systems suggests that spines are evolutionarily ancient evolution and thus could be effectively investigated in simpler model organisms^5, 6^. *C. elegans* offers several advantages for this effort. Because *C. elegans* is transparent, live imaging does not require surgery or other invasive methods that are typically necessary for *in vivo* imaging of spines in an intact mammalian nervous system. Genetic tools in *C. elegans* are also especially well-developed. Of particular importance, are forward genetic screens that could be used to reveal new determinants of spine assembly. A recent study, for example, reported that neurexin, a conserved membrane protein and established regulator of synaptic assembly, is necessary for spine morphogenesis in *C. elegans* DD GABAergic neurons. Interestingly, in this case, neuroligin, the canonical neurexin ligand, is not required, suggesting a potentially new neurexin-dependent mechanism of synaptogenesis^13^.

This study features the first images of spines in motor neurons of an intact, living animal^26^. Our imaging experiments by light and electron microscopy confirmed that both DD and VD motor neurons display dendritic spines (Figure 1 and S3). Thus neurons reported to have “short branches” in the original EM reconstruction of the adult *C. elegans* hermaphrodite nervous system^10^, the cholinergic (RMD, SMD) and GABAergic (RME, VD, DD) motor neurons and the interneuron RIP, are also likely to display *bona fide* dendritic spines. An ongoing effort to produce a gene expression fingerprint of each type of *C. elegans* neurons could be useful for identifying genetic programs that are uniquely correlated with spine morphogenesis since only a small number^9^ of *C. elegans* neurons are reported to have spine-like structures^56^. Finally, the developmentally regulated remodeling program undergone by DD neurons^57–59^ transforms presynaptic boutons into postsynaptic spines in larval DD neurons and thus could be especially informative for live imaging studies of synaptic plasticity and spine morphogenesis.

## ACKOWLEDGEMENTS

We thank M. Francis for sharing unpublished observations. We thank Douglas Holmyard (Nano Nanoscale Biomedical Imaging Facility) for preparing serial EM sections, the laboratory of Dylan Burnette for insightful imaging discussions and Erin Miller for help with aligning serial electron micrographs. Some *C. elegans* strains used in this work were provided by the Caenorhabditis Genetics Center, which is funded by the NIH National Center for Research Resources (NCRR). Super-resolution imaging was acquired at the Vanderbilt Cell Imaging Shared Resource (1S10OD201630-01). This work was supported by National Institutes of Health grants to DMM (R01NS081259 and R01NS106951), and by the Canadian Institute of Health Research grant to MZ (FS154274). ACC is supported by an AHA predoctoral fellowship (18PRE33960581).

## AUTHOR CONTRIBUTIONS

A.C.C. and D.M.M. conceived the project. A.C.C. designed and performed super-resolution and live-imaging, as well as genetic manipulations to test effects on spine density and Ca^++^ transients. A.C.C and B.M. collected, aligned and analyzed serial sections. DM (see acknowledgement) prepared and sectioned the EM sample; B. M. analyzed and 3D reconstructed the EM micrographs. S.H. performed the NATF labeling of LEV-10. S.P. contributed to development of directed Ca^++^ imaging. A.C.C, and D.M.M. wrote the manuscript with input from all the authors.

## METHODS

### Worm Breeding

Worms were maintained at 20°- 25°C using standard techniques^60^. Strains were maintained on NGM plates seeded with *E. coli* (OP-50) unless otherwise stated. The wild type (WT) is N2 and only hermaphrodite worms were used for this study. Staging as L3, early L4 (eL4), L4 and young adult worms was defined following vulva development as previously reported^61^.

### Molecular Biology

Gateway cloning was used to build pACC06 (*punc-25::*LifeAct::GFP). Briefly, plasmid pDONR221 (Plastino Lab)^62^ was used in the LR reaction with pMLH09 (*punc-25*::ccdB::GFP) to create pACC06. Additional plasmids were created using InFusion cloning (Takara). The InFusion cloning module (SnapGene) was used to design primers to create the desired plasmid. Briefly, vector and insert fragments were amplified using CloneAmp HiFi polymerase. PCR products were gel-purified and incubated with In-Fusion enzyme for ligations. Constructs were transformed into Stellar Competent cells and confirmed by sequencing (See full list of plasmids). Plasmids are available upon request. Addgene provided sequences for GCamP6s (#68119), and ceChrimson (#66101)^48^. miniSOG sequence was a gift from the Jin lab^55^. pSH40 was a gift from the Bargmann Lab^54^.

### Feeding RNAi

Clones from the RNAi feeding library (Source BioScience) were used in this study. RNAi plates were produced as described^59^. Briefly, RNAi bacteria were grown in the presence of ampicillin (50 µg/mL) and induced with IPTG (1 mM). 250µL of the RNAi bacterial culture was seeded on NGM plates. RNAi plates were kept at 4°C for up to one week until used. RNAi experiments were set-up as follows: 3 to 5 L4 worms (NC3458) were placed on RNAi plates and maintained at 20°C. Four days later, F1 progeny was imaged as young adults.

### Electron Microscopy

Young adult animals were fixed using high pressure freezing followed by freeze substitution, as previously described^63, 64^, with minor modification: they were held at −90°C in acetone with 0.1% tannic acid and 0.5% glutaraldehyde for 4 days, exchanged with 2% osmium tetroxide in acetone, raised to −20°C over 14h, held at −20°C for 14h, then raised to 4°C over 4h before washing. Additional *en bloc* staining was performed with uranyl acetate for 2h at room temperature, followed by lead acetate at 60°C for 2h. Samples were embedded in Epon, cured at 60°C for 24h, then cut into 50nm-thick serial sections. Sections were not poststained. Images were taken on an FEI Tecnai 20 transmission electron microscope with a Gatan Orius digital camera, at 1nm/pixel.

### 3D reconstruction

Images were aligned into a 3D volume and segmented using TrakEM2^65^, a Fiji plugin^66^. Neuron identity was assigned based on characteristic morphology, process placement, trajectory and connectivity^10, 64^. The ventral and dorsal cord volumes contained the anterior-most 25 um of DD1, and 27 um of VD2, respectively. Volumetric reconstructions were exported to 3Ds Max for processing (3Ds Max, Autodesk)

### AiryScan Microscopy

Worms were mounted on 10% agarose pads and immobilized with 15mM levamisole/0.05% tricaine dissolved in M9. A Zeiss LSM880 microscope equipped with an AiryScan detector and a 63X/1.40 Plan-Apochromat oil objective lens was used to acquire super resolution images of the DD neuron. Images were acquired as a Z-stack (0.19µm/step), spanning the total volume of the DD ventral process and submitted for AiryScan image processing using ZEN software. Developmental stage was determined by scoring gonad and vulva development^67^.

### Classification of spines

Spine shapes were determined from Z-projections of AiryScan images and by 3D-EM reconstruction. Mean and SD were determined using GraphPad. Spines were classified as thin/mushroom, filopodial, stubby or branched. Thin/mushroom spines displayed a constricted base (neck) and an expanded tip (head). Filopodial spines do not have a constricted base (no neck) but are protrusions of constant width. Stubby spines were recognized as protrusions with a wide base and tip. Branched spines were identified as protrusions with more than one visible tip.

### Ribosomal protein labeling in DD spines

To label ribosomes in DD spines, we used DD-specific RIBOS^46^. To label DD spines we injected *Pflp-13::*LifeAct::mCherry plasmid into CZ20132 and used AiryScan imaging to examine transgenic animals (See AiryScan microscopy section).

### Actin dynamics

A Nikon microscope equipped with a Yokogawa CSU-X1 spinning disk head, Andor DU-897 EMCCD camera, high-speed piezo stage motor, 100X/1.49 Apo TIRF oil objective lens and a 1.5X magnification lens was used for live imaging. For measurements of LifeAct::GFP and cytosolic mCherry dynamics, young adults (NC3315 and XMN46) were mounted on 10% agarose pads and immobilized with 15mM levamisole/0.05% tricaine dissolved in M9. Z-stacks (0.5µm/step) were collected every 3 minutes. Movies were submitted to 3D-deconvolution on NIS-Elements using the Automatic algorithm and aligned with the NIS Elements alignment tool. For each movie, ROIs were defined along the dendritic shaft for each spine. Mean ROI Intensity was calculated for each time point and exported to Microsoft Excel. Background was determined from a neighboring region inside the worm and subtracted from the ROI in each timepoint. Mean intensity changes where normalized to the mean Intensity from the first timepoint of each movie. Intensity changes for LifeAct::GFP and mCherry were graphed using Prism6 software.

### GCaMP6s dynamics in DD spines

GCaMP6s imaging was performed on a Nikon microscope equipped with a Yokogawa CSU-X1 spinning disk head, Andor DU-897 EMCCD camera, high-speed piezo, 100X/1.49 Apo TIRF oil objective lens and a 1.5X magnification lens. NC3484 worms were immobilized using a combination of 3µL of 100mM muscimol (TOCRIS biosciences #0289) and 7µL 0.05um polybeads (2.5% solids w/v, Polysciences, Inc. #15913-10). Triggered acquisition was used to excite the GCaMP and mCherry signals with 488nm and 561nm lasers. Single plane movies were collected every second for at least 24 seconds. Movies were submitted for 2D-deconvolution on NIS-Elements using the Automatic algorithm. Movies collected with NIS-elements were aligned through time using the ND alignment tool. ROIs with the same area for each channel were defined in spines and on a neighboring region to determine background intensity for every time point. Mean ROI intensity was exported to Microsoft Excel for subtraction of mean fluorescence background intensity. Fluorescence at each timepoint was normalized to intensity at t=0 for GCaMP6s and mCherry signals. Local peaks of GCaMP6s fluorescence were identified between neighboring neurons and the difference between the timepoints (deltaT) was calculated. Traces were graphed on Prism6.

To detect evoked calcium responses in DD neurons, NC3569 were grown for 1 generation on an OP-50-seeded plate with freshly added ATR or carrier (EtOH). L4 worms were glued (Super Glue, The Gorilla Glue Company) to a microscope slide in 2µL 0.05um polybeads (2.5% solids w/v, Polysciences, Inc. #15913-10) plus 3µL of M9 buffer and imaged under a coverslip. GCaMP6s imaging was performed on a Nikon microscope equipped with a Yokogawa CSU-X1 spinning disk head, Andor DU-897 EMCCD camera, high-speed piezo and 100X/1.49 Apo TIRF oil objective lens. Single plane images encompassing DD1 postsynaptic spines and adjacent VA and DA motor neurons were collected at 2 frames/second for 15 seconds. The sample was illuminated with a 561nm laser at 2.5 sec intervals (e.g., every 5^th^ frame) for red light activation of Chrimson expressed in cholinergic DA and VA motor neurons (*Punc-4:*:ceChrimson::SL2::3xNLS::GFP) while maintaining constant illumination with a 488nm laser to detect GCamP6s signals. For quantifying GCaMP6s fluorescence, videos were aligned and 2D-deconvolved using NIS Elements software. ROIs were drawn on the spines and on a nearby region to capture background fluorescence. Mean fluorescence intensity of each ROI for each frame was exported into Excel for further analysis. Background was subtracted from each frame and measurements were normalized to t = 0. Mean fluorescence traces were plotted using Prism6. Standard deviations of each time point between conditions (EtOH vs ATR) or ceChrimson activation (before vs after) were compared using the F-test in Prism6.

### Temporal silencing with histamine chloride

Gravid adults were allowed to lay eggs for 2 hours on an OP50-seeded plate at 20 C to produce a synchronized population of L1 larvae. The middle time-point of the egg-laying session was considered T_0_. At T_19_ (time in hours), L1 larvae were transferred to control or histamine plates and maintained at 20°C until imaging on an LSM880 AiryScan microscope at the young adult stage. For control plates, 200µL of water was added to OP-50 seeded NGM plates. For histamine plates, 200µL of 0.5M Histamine, diluted in water, was added to OP-50 seeded NGM plates.

### Temporal neuronal activation

Gravid adults were allowed to lay eggs for 2 hours on an OP-50-seeded plate with freshly added ATR. The resultant synchronized population of L1 larvae was maintained at room temperature (23-25C). At T_27_ (L2 larvae), we used WormLab (MBF Bioscience) for exposure to repetitive cycles of 1 second ON + 4 seconds OFF for 6 hours of a 617nm precision LED (Mightex PLS-0617-030-10-S). Images were collected on an LSM880 AiryScan microscope with 60X/1.4 Plan-Apochromat oil objective lens. Control worms were not exposed to light but grown on ATR plates. 100mM ATR (Sigma, #A7410) was prepared in ethanol and stored at −20C. 300uM of ATR was added to OP-50 bacteria and seeded on NGM plates. Plates were dried in darkness overnight and used the next day for experiments.

### Image Analysis

FIJI^66^ and NIS-Elements software were used for data quantification (Figure 1E-J, 2D, 2G, 2J, 4C, 4D, 4G, 4H, 5E, 5I and 5M). Z-stacks were flattened in a 2D projection and line scans were manually drawn along protrusions and perpendicular to the proximal shaft (Figure 1d) to determine the Protrusion/Shaft ratio (Figure 1e). Spine density was calculated using the counting tool in NIS Elements and then normalized to number of spines per 10 µm of dendrite length.

### Statistical Analysis

For comparison between 2 groups, Student’s T-test was used and p<0.05 was considered significant. ANOVA was used to compare between 3 or more groups followed by Dunnett’s multiple-comparison test. Standard Deviations between two samples were compared using an F-test and considered p<0.05 as significant.

### Ablation of DD neurons

Twenty gravid adults (NC3480) were allowed to lay eggs for 2 hours with the middle time point considered T_0_. At T_16_, (hours) DD neurons were ablated by miniSOG^55^ activation by exposing worms for 45 min to a 470nm LED light (#M470L2, Thor Labs). Animals were then maintained at 20°C until imaging at ∼T_60_ as young adults.

