## Supplementary Information for "Dendritic spines on GABAergic neurons respond to cholinergic signaling in the *Caenorhabditis elegans* motor circuit"

**Figure S1. Dendritic spines adopt distinct morphologies and display a dynamic actin cytoskeleton.**

**A.** Spine densities determined from young adult DD neurons labeled with mCherry (3.77 ± 0.9 spines/10μm) vs LifeAct::GFP (4.36 ± 1.0 spines/10μm) are not significantly different and are comparable to spine density determined by 3D EM reconstruction of DD1 (4.2 spines/10μm) (dashed blue line). T-test between mCherry and LifeAct::GFP $p>0.05$, $n>16$. Data for mCherry and LifeAct::GFP also appears in Figure 1F.

**B.** Frequency of spines by type: Thin/Mushroom (55.54 ± 14.54%), Filopodial (10.27 ± 8.70%), Stubby (18.78 ± 10.74%), Branched (15.42 ± 6.01%). Dashed blue lines denote frequency for each spine type from EM reconstruction of DD1: Thin/Mushroom (58.3%), Filopodial (25%), Branched (8.3%) and Stubby (8.3%).

**C-G.** Live-imaging of **(C,D)** LifeAct::GFP and **(E,F)** mCherry-labeled DD spines of L4 stage larvae reveals dynamic LifeAct::GFP vs stable cytosolic mCherry signals**. (D,F)** Normalized traces of LifeAct::GFP and mCherry fluorescence from live imaging. Different shades of green and red lines are used to allow visualization of individual traces. **(G)** Comparison of standard deviations of all traces for each timepoint reveals significantly different variance between LifeAct::GFP vs cytosolic mCherry markers (F-test, * is $p < 0.0001$ from $T_3$ to $T_{30}$ and $ is $p < 0.05$ at $T_{39}$, $n > 51$ traces). Scale bars = 500 nm.

**H.** Regulators of actin polymerization define spine density. (Left) Representative images (young adults) of DD spines for WT (wild type), *toca-1 (tm2056)*, empty vector (control), *wave/wve-1*(component of WRC) RNAi and *p21/arx-5* (component of Arp2/3 complex)

RNAi**.** (Right) WT young adults show 4.36 ± 1.0 spines/10 µm vs *toca-1* (*tm2056*) with 2.77 ± 1.1 spines/10 µm (t test, *** is $p<0.001$). Spine density is reduced with RNAi of *wave/wve-1* (2.24 ± 0.7 spines/10 µm) or *p21*/*arx-5* RNAi (2.20 ± 0.7 spines/10 µm) versus Empty Vector (E.V.) control (2.9 ± 0.4 spines/10µm). One-way ANOVA with Dunnett's multiple comparison test, ** is $p<0.01$, n > 9 worms. Measurements are Mean ± SD. Scale bar is 1 µm.

**Figure S2. Synaptic activity regulates postsynaptic DD spine density.**

**A.** Spine density (spines/10 µm) increases throughout development in the wild type (WT). L3 (2.21 ± 0.7), (early L4) eL4 (2.62± 0.8), L4 (3.39 ± 0.9) and (Young Adult) YA (3.92 ± 1.01). One-way ANOVA of all groups vs L3 shows that spine densities at L4 and YA stages are different from L3. ** is $p<0.01$ and **** is $p<0.0001$. Data for YA are the same as in Figure S1H for WT.

**B.** Spine density (spines/10 um) does not increase during development with reduced cholinergic signaling in *unc-17(e113)*. L3 (2.91 ± 1.0), eL4 (2.76 ± 0.9), L4 (2.32 ± 0.87) and YA (2.84 ± 0.9). One-way ANOVA of all groups against L3 stage shows no statistically significant difference during development.

**C.** Spine density (spines/10µm) is elevated throughout development in acetylcholinesterase deficient *ace-1(p100);ace-2(g72)* mutant animals. L3 (3.83 ± 1.2), eL4 (4.17 ± 0.8), L4 (3.16 ± 1.2) and YA (3.49 ± 1.1). One-way ANOVA of all groups against L3 stage shows no statistically significant difference during development.

**D.** Spine density (spines/10µm) does not increase during development in *unc-31(e169)* mutants with impaired dense core vesicle release L3 (2.18 ± 1.0), eL4 (2.45 ± 1.1), L4

(2.65 ± 0.9) and YA (1.89 ± 0.8). One-way ANOVA of all groups against L3 stage shows no statistically significant difference during development.

**E.** Spine density (spines/10µm) does not increase during development in *unc-31(e169)* and shows lower spine density than WT at the Young Adult (YA) stage. T-test at each timepoint between genotypes, **** is p<0.0001. Measurements are mean ± SD.

**Figure S3. Ventral D-GABAergic motor neurons have dendritic spines**

**A.** Thirteen VD motor neurons (green) are located in the ventral nerve cord. In the adult, VD presynaptic boutons (oblong ovals) innervate ventral muscles (gray cells) and VD postsynaptic termini (spines) receive cholinergic input from DA and DB motor neurons (purple) on the dorsal side.

**B-D.** GABAergic VD neurons show dendritic spines. GABA neurons were labeled with LifeAct::GFP (*Punc-25::LifeAct::GFP*) and DD neurons expressed miniSOG (*Pflp-13::miniSOG::SL2::BFP*). **(B)** Synchronized L1 larvae were exposed to blue light for 45 minutes during $T_{16}$-$T_{17}$ (hours after egg laying) to ablate DD neurons and maintained at 20°C until the adult stage. **(C)** AiryScan imaging of the dorsal cord in young adults ($T_{60}$) shows spine-like protrusions on VD neurons. **C'.** Zoomed in region of C. Scale bars = 1 µm. **(D)** VD spine density (3.007 ± 1.43 spines/10µm), n = 17. Dashed blue line represents spine density detected in VD2 with 3D EM (3.33 spines/10µm).

**E.** 3D EM reconstruction shows mitochondria in the shaft of VD dendrites (arrows) but not in VD spines (clear arrowheads).

**F.** 3D EM reconstruction shows that 88.89% (n = 8/9) of VD spines directly appose presynaptic dense projections from either DA, DB or AS motor neurons (black and clear arrowheads). 11.11% (n = 1/9) of VD spines do not oppose a presynaptic terminal (gray arrowhead). 50% (n = 4/8) of VD spines are contacted by a single synaptic partner (black arrowheads) whereas 50% (n = 4/8) of VD spines are contacted by more than one partner (clear arrowheads).

**Video S1-S2** Live-imaging of cytosolic mCherry in DD spines.

**Video S3-S4** Live-imaging of LifeAct::GFP in DD spines.

**Video S5** Spine contacted by two presynaptic partners (EM reconstruction)

**Video S6** Serial EM reconstruction of DD1.

**Video S7** Endogenous $Ca^{++}$ waves in DD spines, cytosolic mCherry.

**Video S8** Endogenous $Ca^{++}$ waves in DD spines, GCaMP6s.

**Video S9** Evoked $Ca^{++}$ waves in DD1 spine treated with ATR.

**Video S10** No evoked $Ca^{++}$ waves in DD1 spine treated with EtOH.

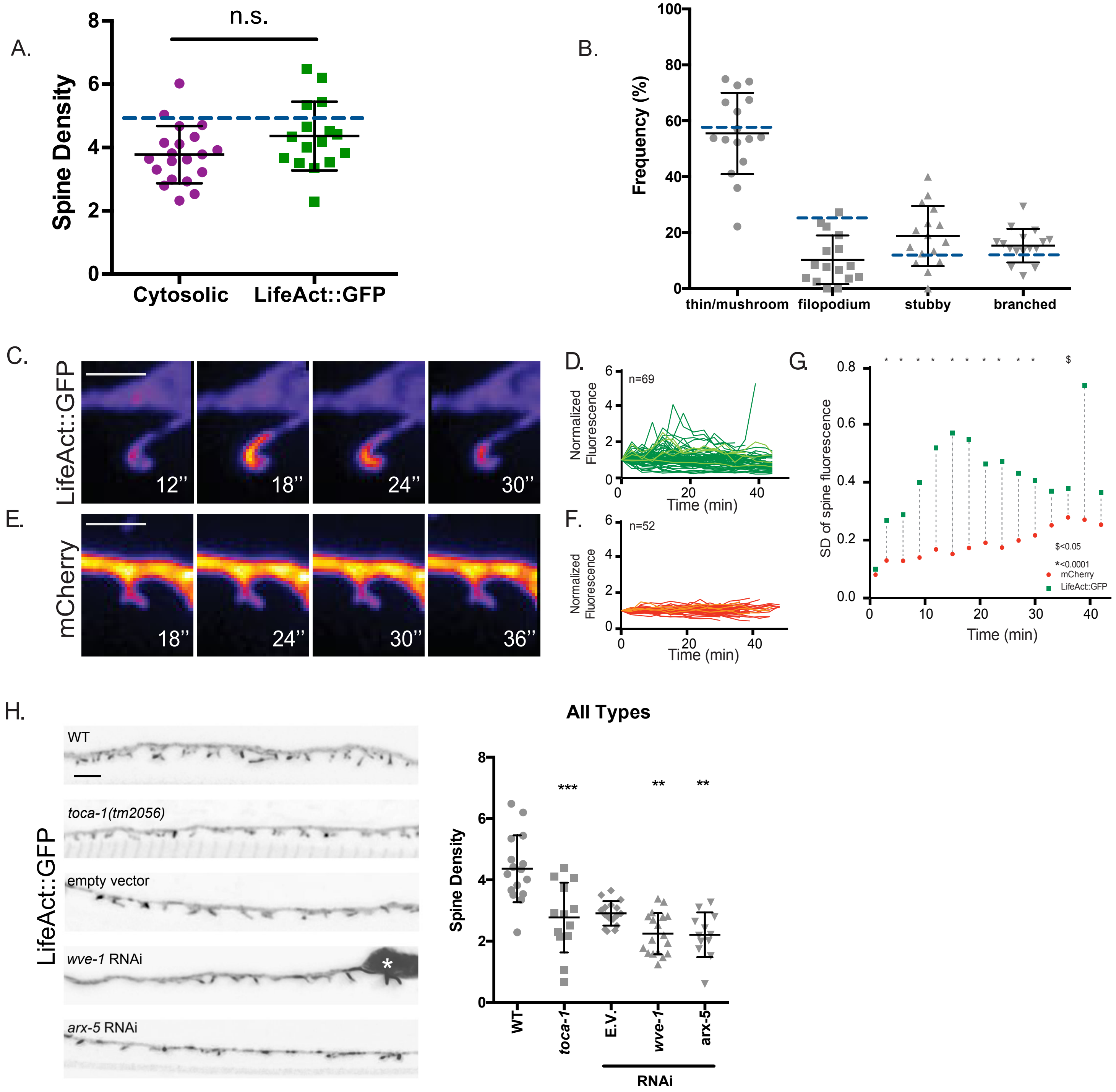

A.
n.s.
Spine Density
8
6
4
2
0
Cytosolic
LifeAct::GFP
B.
Frequency (%)
100
80
60
40
20
0
thin/mushroom
filopodium
stubby
branched
C.
LifeAct::GFP
12''
18''
24''
30''
D.
Normalized Fluorescence
n=69
Time (min)
E.
mCherry
18''
24''
30''
36''
F.
Normalized Fluorescence
n=52
Time (min)
G.
SD of spine fluorescence
Time (min)
$<0.05
*<0.0001
mCherry
LifeAct::GFP
H.
WT
toca-1(tm2056)
empty vector
wve-1 RNAi
arx-5 RNAi
LifeAct::GFP
All Types
Spine Density
***
**
**
WT
toca-1
E.V.
wve-1
arx-5
RNAi

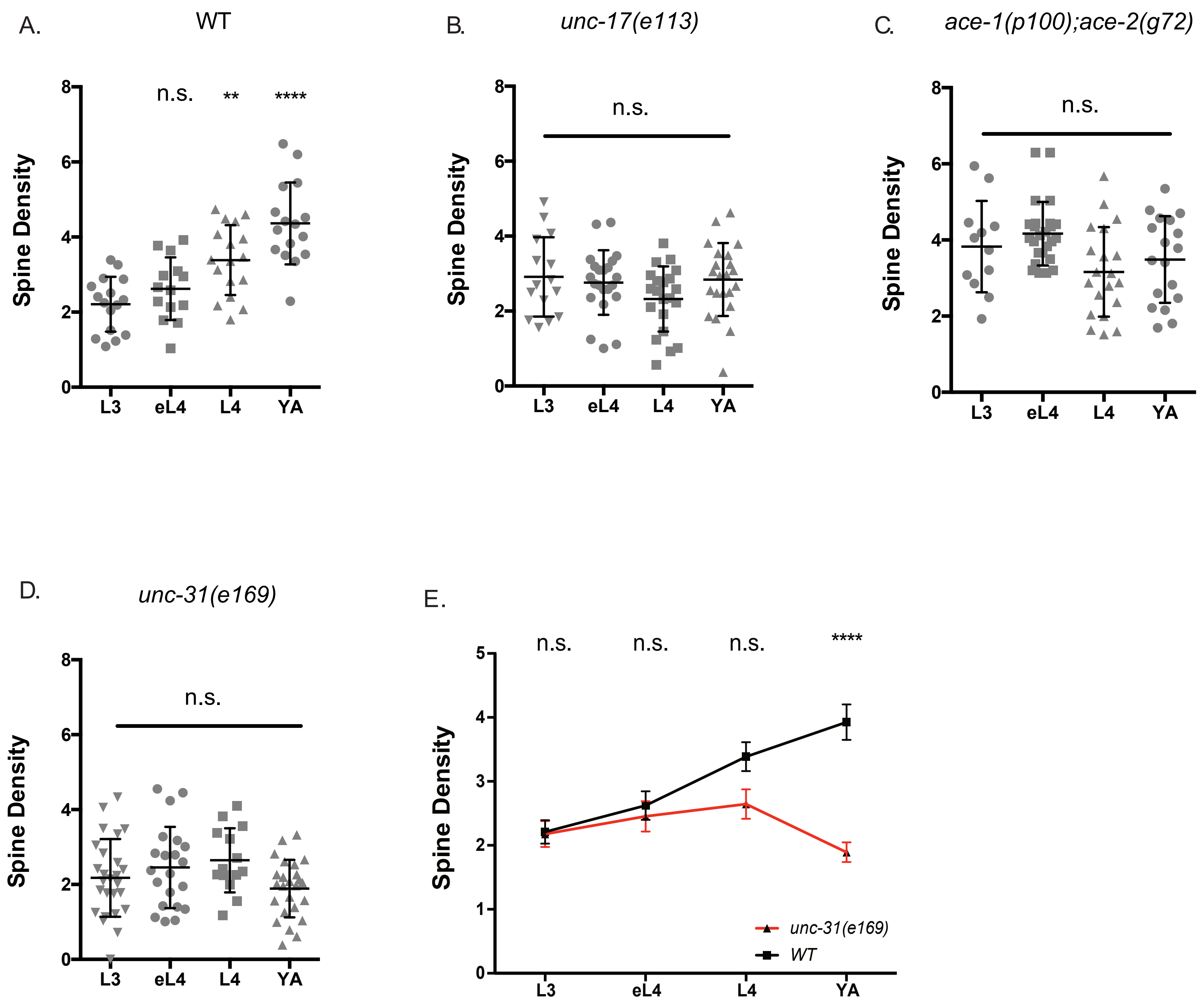

A.
WT
Spine Density
n.s.
**
****
L3
eL4
L4
YA
B.
unc-17(e113)
n.s.
C.
ace-1(p100);ace-2(g72)
D.
unc-31(e169)
E.
unc-31(e169)
WT

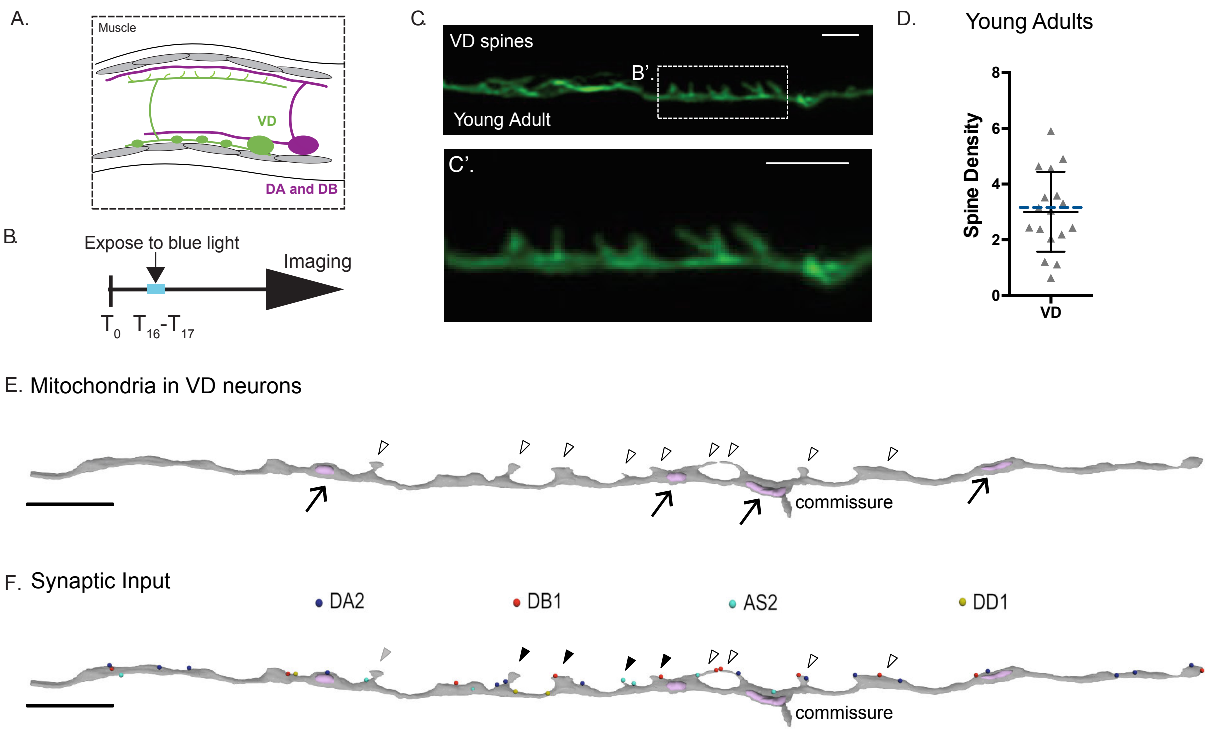

A.
Muscle
VD
DA and DB
B.
Expose to blue light
Imaging
T0 T16-T17
C.
VD spines
B'.
Young Adult
C'.
D.
Young Adults
Spine Density
8
6
4
2
0
VD
E. Mitochondria in VD neurons
commissure
F. Synaptic Input
DA2
DB1
AS2
DD1
commissure
